# Smooth muscle LRRC8A knockout preserves vascular function in AngII hypertension

**DOI:** 10.1101/2025.05.08.652978

**Authors:** Hyehun Choi, Sourav Panja, Hong-Ngan Nguyen, Ryan J. Stark, Fred S. Lamb

## Abstract

Angiotensin II (AngII) causes hypertension and vascular inflammation both directly and indirectly via cytokines. In vascular smooth muscle cells (VSMCs) AngII and TNFα activate NADPH oxidase 1 (Nox1) to produce superoxide. TNFα receptors associate with Nox1 and Leucine Rich Repeat Containing 8A (LRRC8A) anion channels to modulate inflammation, as well as contractility in a RhoA-dependent manner. VSMC-specific LRRC8A knockout (KO) mesenteric arteries are protected from TNFα-induced injury and vasodilated better. We hypothesized that LRRC8A KO would preserve vascular function and decrease blood pressure (BP) in AngII-infused mice. Wild type (WT) and LRRC8A KO mice received AngII infusions for 14 days. Systolic BP was not different, but KO mice had more BP “dipping” during inactive periods and dipping was preserved after AngII. Contraction of KO mesenteric vessels to AngII itself was not altered, however after AngII exposure the function of KO aortic and mesenteric vessels was less impaired as reflected by less augmented contraction to norepinephrine and serotonin and preserved relaxation to acetylcholine and sodium nitroprusside. Western blotting revealed increased soluble guanylate cyclase alpha and reduced CPI-17 in hypertensive KO aortae. Consistent with the presence of lower Rho kinase activity in KO VSMCs, phosphorylation of Ezrin/Radixin/Moesin (ERM) and Cofilin was reduced. Aortic wall/lumen was not different between hypertensive WT and KO mice, but AngII caused less proliferation (lower PCNA), and less induction of antioxidant enzymes and senescence markers in KO vessels. Thus, while AngII-induced contraction does not require LRRC8A, these channels support the associated inflammatory response which modifies BP dipping.

## Introduction

Chronic vascular inflammation is a critical driver of hypertension^1^, impacting both structure and function^2^. The signaling pathways of many key cytokines and vasoactive mediators share a requirement for extracellular superoxide (O_2_^•-^) production via activation of an NADPH oxidase (Nox)^3^. Frequently there is also an associated increase in mitochondrial reactive oxygen species (ROS) production that potentiates the oxidative stress^1^ that is a fundamental characteristic of the hypertensive state^4^. This inflamed environment promotes the premature development of senescence, an aging-related process in which cells stop proliferating and become dysfunctional^5,6^.

Angiotensin II (AngII) is a powerful vasoconstrictor and pro-inflammatory agent that is closely linked to the pathogenesis of hypertension. It drives changes in vascular structure and vasomotor function that underly the risk of life-threatening complications of cardiovascular disease such as myocardial infarction and stroke. Therefore, antihypertensive agents frequently target key aspects of this pathway. In vascular smooth muscle cells (VSMCs) AngII receptors activate Nox1, producing extracellular O_2_^•-^ and other ROS^7^ that promote oxidative stress and vascular dysfunction^8–10^. AngII also triggers other pro-inflammatory pathways including tumor necrosis factor α (TNFα) signaling. Mice lacking this cytokine are protected from AngII-induced hypertension and high blood pressure (BP) is restored by concurrent TNFα infusion^11^. TNFα can independently enhance vascular contractility and cause endothelial dysfunction^12^, and VSMC-specific TNFα knockout or treatment with the TNFα blocker Etanercept reduces vascular myogenic tone and BP^13^.

TNFα signaling and related VSMC inflammation are attenuated by Nox1 knockdown^14,15^ and Leucine Rich Repeat Containing 8A (LRRC8A), which is a critical component of volume-regulated anion channels, is part of a Nox1-containing multi-protein complex^12^. Decreasing LRRC8A expression or pharmacologic block of these channels abrogates TNFα-induced Nox1 activation, O_2_^•-^ production, MAPK singling and vascular dysfunction^12,16^. AngII increases LRRC8A expression in cerebral blood vessels^17^ and loss of these channels decreases systolic BP (SBP) and abrogates AngII-induced cerebrovascular remodeling^18,19^. We therefore hypothesized that LRRC8A is required for the AngII-induced vascular inflammation and changes in vasomotor function that are thought to contribute to the development of hypertension in this model.

VSMC-specific LRRC8A knockout (KO) mice were exposed to a 2-week subcutaneous infusion of AngII. Importantly, lack of LRRC8A protein did not alter the contractile response of mesenteric resistance vessels to AngII *in vitro*, and the BP of AngII-infused mice was only significantly reduced during light cycles when these nocturnal animals were less active or sleeping (preserved BP dipping). Despite these relatively small differences in BP, the responses of isolated KO blood vessels to vasoconstrictors (norepinephrine, serotonin) and both endothelium-dependent (acetylcholine) and independent (sodium nitroprusside) vasodilators were remarkably protected from the AngII-induced dysfunction seen in wild type (WT) mice.

KO vessels also displayed reduced phosphorylation of protein phosphatase 1 regulatory subunit 14A (PPP1R14A), a.k.a. protein kinase C-potentiated inhibitor protein of 17 kDa (CPI-17), as well as ERM proteins (Ezrin/Radixin/Moesin) and cofilin all of which are regulated by Rho kinase (ROCK). This supports our prior proposal that reduced ROCK activation in LRRC8A KO VSMCs limits contractility and improves vasodilation^12^. Antioxidant enzymes (catalase, heme oxygenase-1 (HO-1)), an inflammation-associated protein (iNOS; inducible nitric oxide synthase) and a marker of proliferation (PCNA; proliferating cell nuclear antigen) were all significantly reduced in LRRC8A KO AngII-exposed vessels compared to WT. KO aortae also expressed lower levels of β-galactosidase, consistent with less vascular senescence. Collectively, these data suggest that reduced inflammation and lower oxidative stress in AngII exposed VSMCs lacking LRRC8A channels protects vascular function that manifests as preservation of BP dipping with inactivity.

## Methods

### Animals

All procedures were performed in accordance with the *Guiding Principles in the Care and Use of Animals*, approved by the Vanderbilt University Institutional Animal Care and Use Committee. VSMC-specific LRRC8A KO mice were created by crossing floxed LRRC8A mice provided by Dr. Rajan Sah (Washington University, St. Louis, MO)^20^ with SM22α-Cre mice (Jackson Laboratory, Bar Harbor, ME). Genotyping was performed by TransnetYX, Inc. (Memphis, TN) using PCR analysis of tail-tip DNA. WT and KO littermates underwent experimentation. The mice were housed on a 12-hour light/dark cycle, fed a standard chow diet, and provided with water *ad libitum*.

### Blood pressure measurement

Blood pressure was measured via telemetry (transducer Model TA11PAC-10, Data Sciences International, St. Paul, MN) by the Vanderbilt Mouse Metabolic Phenotyping Center. For implantation, mice (25-30g) were given analgesia using Ketoprofen (Ketofen Injectable Solution, Zoetis Inc, Kalamazoo, MI) at a dose of 5mg/kg and anesthetized using Isoflurane Inhalant (ISOFLURANE USP, Liquid for Inhalation, PIRAMAL Pharma Limited, Telanggana State, India). A SomnoFlo Vaporizer (Kent Scientific, Torrington, CT) delivered the inhalant anesthetic at an induction rate of 350mL/min with 2.5% isoflurane, and a maintenance flow rate of 100-150 mL/min with 1-1.5% isoflurane. After isolation of the left common carotid artery, the catheter connecting transducer was introduced into the carotid artery and advanced until the tip was inside the thoracic aorta. The transducer was positioned subcutaneously along the left flank near the hindlimb. Mice were allowed to recover from the surgery for 1 week and baseline BP was then recorded for 2 days. An osmotic minipump (Durect Corporation, Cupertino, CA) with AngII (1 µg/kg per min, Phoenix Pharmaceuticals, Burlingame, CA) was implanted subcutaneously into the dorsum and BP was monitored for an additional 14 days. Data was recorded using the DSI PONEMAH software.

### Aortic and mesenteric vascular reactivity

WT and KO littermates were randomly chosen to be sham or AngII groups. AngII (3 µg/kg per min) was delivered using osmotic minipumps implanted subcutaneously for 14 days and sham groups were exposed to the same surgical procedures. Mice were euthanized by CO_2_ inhalation/cervical dislocation and blood vessels were harvested. Hypertension was confirmed by measurement of heart to body weight ratio which increased in all AngII-treated mice, and to a similar degree in the WT and KO groups (Suppl Table S1). Thoracic aortae and mesenteric arteries (∼100 µM inner diameter, first-or second-order branches of superior mesenteric artery) were excised, cleaned of associated fat and cut into 2 mm (aorta) or 1-2 mm (mesenteric) length-rings in ice-cold physiological salt solution (PSS; 130 mmol/L NaCl, 4.7 mmol/L KCl, 1.18 mmol/L KH_2_PO_4_, 1.18 mmol/L MgSO_4_·7H_2_O, 1.56 mmol/L CaCl_2_·2H_2_O, 14.9 mmol/L NaHCO_3_, 5.6 mmol/L glucose, and 0.03 mmol/L EDTA). The rings were mounted in wire myographs (Danish Myo Technology A/S, Aarhus, Denmark) containing warmed (37 °C), oxygenated (95% O_2_/5% CO_2_) PSS and equilibrated for 1 hour (aorta) or 45 min (mesenteric arteries) under a passive force of 5 mN or 2-2.5 mN respectively. Arterial viability was assessed by stimulation with 120 mmol/L KCl. After washing, rings were stimulated with phenylephrine (PE, 10^-6^ mol/L), followed by exposure to acetylcholine (ACh, 10^-5^ mol/L). A more than 70% relaxation response to ACh was taken as evidence of an intact endothelial layer. AngII-induced contraction study was performed in WT mesenteric arteries and each ring was exposed to only one concentration of AngII. Contractile responses were assessed by cumulative exposure to serotonin (5-HT, 10^-9^ to 10^-5^ mol/L) or norepinephrine (NE, 10^-9^ to 10^-5^ mol/L). Endothelium-dependent relaxation was assessed following PE-induced contraction (1-2 x 10^-6^ mol/L, ∼ ED_75_ dosing) by cumulative addition of ACh (10^-9^ to 3×10^-5^ mol/L), while endothelium-independent relaxation was similarly tested using sodium nitroprusside (SNP, 10^-9^ to 10^-5^ mol/L). Contractile responses were recorded as changes in tension (mN) from baseline, expressed as a percentage of the response to 120mM KCl. Relaxation was expressed as a percentage of the stable contraction produced by PE in each ring immediately prior to the first dose of vasodilator. Return to the basal tension recorded before addition of PE was considered as 100% relaxation.

### Cell culture

Primary mesenteric smooth muscle cells were isolated from the first to third branches of mesenteric arteries in male WT or LRRC8A KO mice by explantation. Briefly, the arteries were cleaned from adherent fat in ice-cold PSS. Blood and endothelial cells were removed by passing a wire through the lumen. The vessels were incised longitudinally, placed in culture and maintained in Dulbecco’s Modified Eagle Medium (DMEM) containing 30% fetal bovine serum (FBS) and 1% penicillin/streptomycin in a humidified incubator at 37 °C, 5% CO_2_ atmosphere. After 1 week when cells had migrated out of the tissue fragments the explants were removed and the migrated cells maintained in DMEM supplemented with 10% FBS, 1% penicillin/streptomycin, 1× minimum essential medium non-essential amino acids, 1× vitamins, and 20 mM HEPES.

### Western blot analysis

Isolated abdominal aortae were cleaned of adherent tissues and fat in ice-cold PSS, flash-frozen in liquid nitrogen and stored at-80 °C. Frozen tissues were ground in a liquid nitrogen-chilled mortar and pestle. Proteins from the frozen tissues or isolated mesenteric VSMC were extracted in phosphoprotecting lysis buffer (PPLB; 50 mM Tris-base, 150 mM NaCl, 1 mM EDTA, 1 mM DTT, 10% glycerol, 1% Triton X-100, 0.1% Na-DOC, 0.1% SDS, 10 mM β-glycerophosphate, 20 mM para-nitrophenyl phosphate, 2 mM sodium pyrophosphate, 1 mM Na3VO4, 5 mM NaF, 10 µg/ml aprotinin, and 1 mM phenyl-methylsulfonyl fluoride (PMSF) at pH 7.4) for 1 hour in ice, and then centrifuged for 20 min at 20000g. Supernatants were collected and 30 µg of protein extracts were separated by electrophoresis on a polyacrylamide gel and transferred to nitrocellulose membranes. The membranes were blocked with LI-COR blocking buffer for 1 hour at room temperature and then incubated with primary antibodies in Tris-buffered saline solution with Tween-20 (0.1%) overnight at 4 °C. Antibodies were as follows: sGCα (#12605-1-AP), sGCβ (#19011-1-AP), cofilin (#66057-1-Ig) from Proteintech (Rosemont, IL), p-CPI-17 (#AB52174, Abcam, Cambridge, UK), CPI-17 (#sc-48406), ERM (#sc-271456), GAPDH (#sc-59540) from Santa Cruz Biotechnology (Dallas, TX), p-ERM (#3726), p-cofilin (#3313), Catalase (#14097), HO-1 (43966) from Cell Signaling Technology (Danvers, MA), CuZnSOD (#AF3418), ecSOD (#AF4817) from R&D systems (Minneapolis, MN), iNOS (#610328, BD Transduction, San Jose, CA), LRRC8A (#A304-175A, Bethy Laboratories, Montgomery, TX), PCNA (#05-347, MilliporeSigma, Burlington, MA), β-galactosidase (#GTX134513, GeneTex, Irvine, CA), Tubulin (Vanderbilt Antibody Core, Nashville, TN). Fluorescent secondary antibodies were incubated for 2 hours at room temperature, and signals were quantified using the Odyssey Imaging System and Image Studio software (LI-COR Bioscience, Lincoln, NE).

### Aorta sectioning and histology

Thoracic aortae were isolated from AngII-infused male WT and LRRC8A KO mice, fixed in 10% formalin and embedded in paraffin. The paraffin-embedded aortae were cut into 5-µm-thick sections, deparaffinized with xylene and rehydrated with graded alcohol. These sections were stained with haematoxylin and eosin (H&E) or Picro-Sirius Red (Polysciences, Warrington, PA) to identify collagen. Media and lumen areas were estimated by measuring the perimeter length of the borders between the media and the external and internal elastic laminae. This was used to estimate the circumference of the inner and outer borders of the media. These circumferences were used to obtain radii using the formula, circumference = 2πr. Areas were then estimated using the formula, area = πr^2^. The media-to-lumen ratio was calculated based on the ratio of these measured areas.

### Statistical analysis

Data are expressed as mean ± standard error of the mean (SEM), and ‘n’ represents the number of animals used in the experiments, or of individual experiments in cultured cells.

Concentration-response curves were fitted using a nonlinear interactive fitting program (GraphPad Prism 10, San Diego, CA), and two pharmacological parameters were obtained: the maximal effect generated by the agonist (or E_max_) and EC_50_ (molar concentration of agonist producing 50% of the maximum response). Statistical differences were calculated by unpaired *t*-test, one sample *t*-test or one-way ANOVA. Post hoc testing was performed using Tukey analysis to compare all groups. A probability value less than 0.05 (*p* < 0.05) was considered to be statistically significant.

## Results

### AngII-induced contraction of mesenteric arteries

LRRC8A channels modify Nox1 activity and VSMC inflammation^16,21^, and also impact contractility by changing myosin light chain phosphatase (MLCP) activity^12^. In the absence of an inflammatory stimulus vasodilation of isolated mesenteric vessels to ACh and SNP was enhanced, but contractile responses to KCl and phenylephrine did not differ from WT^12^. Unlike KCl and PE, AngII contraction has been directly linked to Nox1 activation^22,23^ including effects on multiple aspects of calcium-dependent contractile signaling^24^. We evaluated the contractile response of mesenteric arteries to AngII and these responses did not differ between WT and LRRC8A KO (Fig. 1A). The mechanism of AngII-induced contraction was also not fundamentally altered in KO vessels. Transient receptor potential canonical channel TRPC3 and TRPC6 are redox-sensitive and can support Ca^2+^ influx in VSMC^25^. Selective inhibitors of TRPC3 (Pyr3) and TRPC6 (SAR7334) were used to compare their contributions to contraction to 10^-7^M AngII in WT vs. KO mesenteric vessels. Pyr3 reduced contraction to AngII similarly in WT and KO, while SAR7334 did not significantly decrease AngII responses (Fig. 1B).

**Figure 1.**
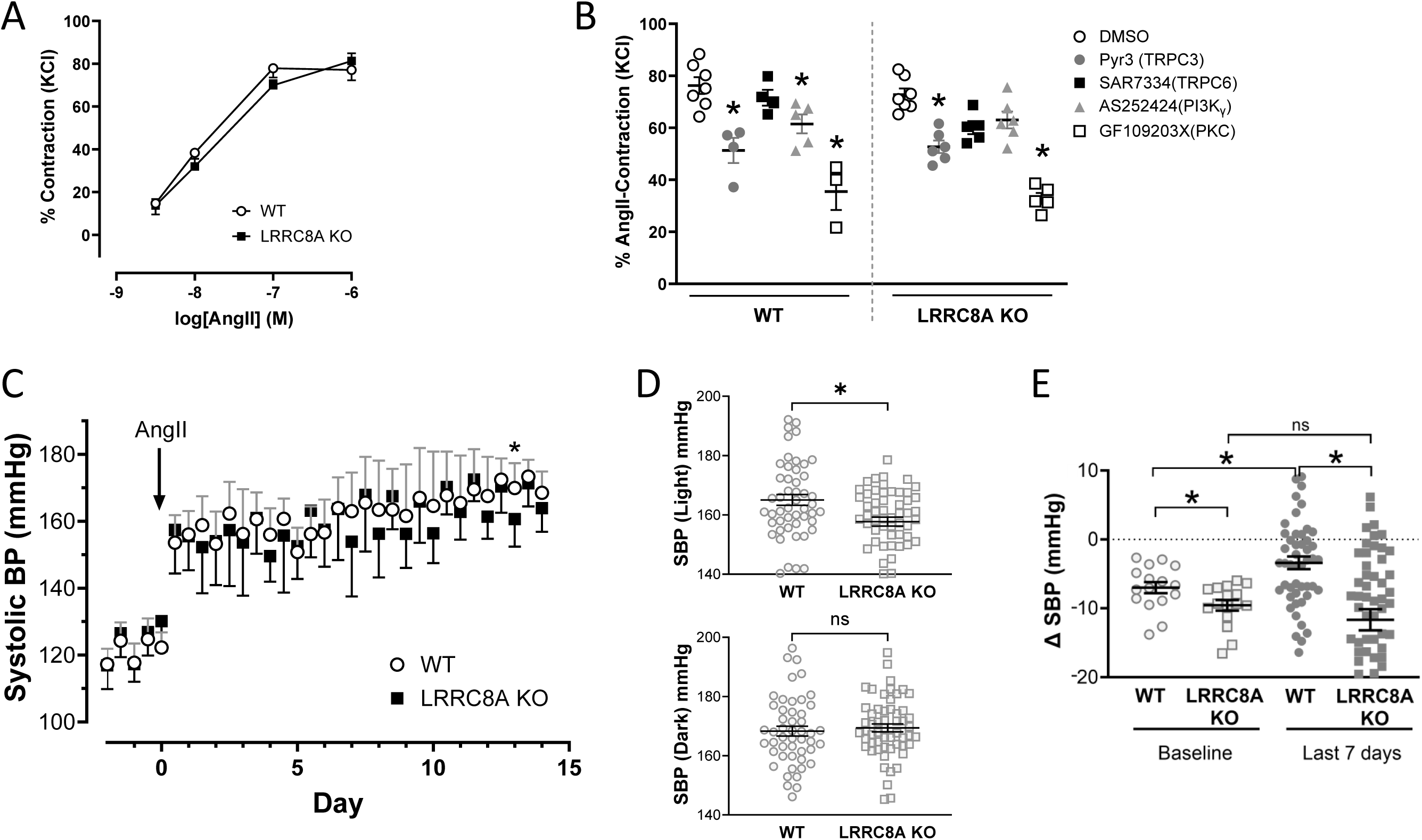
AngII-induced contraction is not altered in mesenteric arteries, but blood pressure dipping is enhanced in LRRC8A KO mice. (**A**) Contraction to AngII was not different in WT and LRRC8A KO vessels (n=5-6). (**B**) Effect of inhibitors of TRPC3 (Pyr3, 5 µM), TRPC6 (SAR7334, 100 nM), PI3Kγ (AS252424, 1 µM) or PKC (GF109203X, 5 µM) (30 min incubation) on contractile responses to AngII (10^-7^ M). * Indicates *p* < 0.05 compared to vehicle (DMSO, n=4-7). (**C**) Systolic blood pressure (SBP) averaged over 12-hour light or dark periods for a 2-day baseline and for 14 days of AngII infusion. (**D**) Blood pressure averages for 12hr periods over the last 7 days of AngII infusion were lower in LRRC8A KO compared to WT when the room was lighted. (**E**) Differences in average SBP between dark and light periods for individual mice during 2-day baseline and over the last 7 days of AngII. * Indicates *p* < 0.05 for WT vs. KO or indicated bar (n=7-8).

Phosphoinositide 3-kinase (PI3K) is activated by AngII and an inhibitor of PI3K_γ_ (AS252424) impaired AngII-induced increases in cytosolic Ca^2+^ concentration and ROS generation^24^.

AS252424 significantly attenuated AngII contraction in WT but not KO vessels, however, the magnitude of contraction did not differ when WT and KO responses were compared directly (Fig. 1B). AngII activates Protein kinase C (PKC) which activates Nox1 and contraction^24^. PKC inhibition with GF109203X impaired AngII contraction to a similar degree in WT and KO (Fig. 1B). Thus, the acute contractile response of mesenteric arteries to AngII does not differ significantly in magnitude or in mechanism between WT and LRRC8A KO mice.

### AngII-induced hypertension

A substantial reduction in tail-cuff SBP was reported in VSMC-specific LRRC8A null mice, both at rest and as early as 1 week after initiation of AngII infusion^18^. Unstressed male WT and LRRC8A KO mice underwent continuous BP monitoring for 2 days and SBP (Fig. 1C and E) and heart rate (HR, Suppl. Fig. S1) did not differ. However, the difference in SBP between active (dark) and inactive (light) periods was significantly larger in KO mice (Fig. 1E, Baseline WT-7.0 ± 0.8 mmHg vs. KO-9.6 ± 0.8 mmHg, \**p*<0.05). Both genotypes responded with a similar brisk rise in SBP in response to AngII but after hypertension was established this difference, termed “dipping”, reduced significantly in WT but was maintained in KO mice (Fig. 1C and E). SBP was lower in KO mice during light but not dark periods (Fig. 1D, SBP (Light) WT 164 ± 1.4 mmHg vs. KO 158 ± 1.0 mmHg for the last 7 days, \**p*<0.05). This was associated with significantly reduced dipping in WT (-3.4 ± 0.9 mmHg), while dipping was preserved in KO mice (-11.7 ± 1.5 mmHg). A similar time-dependent drop HR was also observed in KO mice, but this difference was independent of activity (Suppl. Fig. S1).

### Vascular function in AngII hypertension

AngII-induced hypertension causes changes in vascular responsiveness to both vasoconstrictors and vasodilators that in many cases are Nox1-dependent^26^. Mesenteric arteries and thoracic aorta from WT and LRRC8A KO mice were studied using wire myography.

Previously, mesenteric rings from WT and KO displayed indistinguishable contractile responses to KCl and PE under control conditions^12^. Serotonin (5-HT)-induced vasoconstriction is often markedly altered in hypertension^27^. 5-HT-dependent contraction was decreased in mesenteric arteries from female but not male KO vessels (Fig. 2A and D, E_max_, female WT Sham 144 ± 12.2 vs. KO Sham 87 ± 6.9%, \**p*<0.05). AngII enhanced contraction to 5-HT in both sexes and these responses were again significantly smaller in rings from female KO vessels (Fig. 2A, E_max_, WT AngII 272 ± 15.2 vs. KO AngII 224 ± 12.5%, \**p*<0.05). The 5-HT response of male mesenteric arteries was also slightly more enhanced by hypertension in WT mice (Fig. 2D). Following contraction to PE, KO rings were significantly more sensitive to endothelium-dependent relaxation induced by ACh in both the male and female Sham groups. Consistent with the induction of endothelial dysfunction, the response of AngII hypertensive vessels to ACh was impaired in both WT and KO vessels, but AngII-treated KO rings still responded significantly better than did WT rings (Fig. 2B and E, E_max_, female WT AngII 49.9 ± 4.8 vs. KO AngII 73.5 ± 3.6; male WT AngII 50.6 ± 7.1 vs. KO AngII 74.5 ± 4.6%, \**p*<0.05). Endothelium-independent relaxation to SNP directly assesses the response of the smooth muscle to nitric oxide (NO^•^).

**Figure 2.**
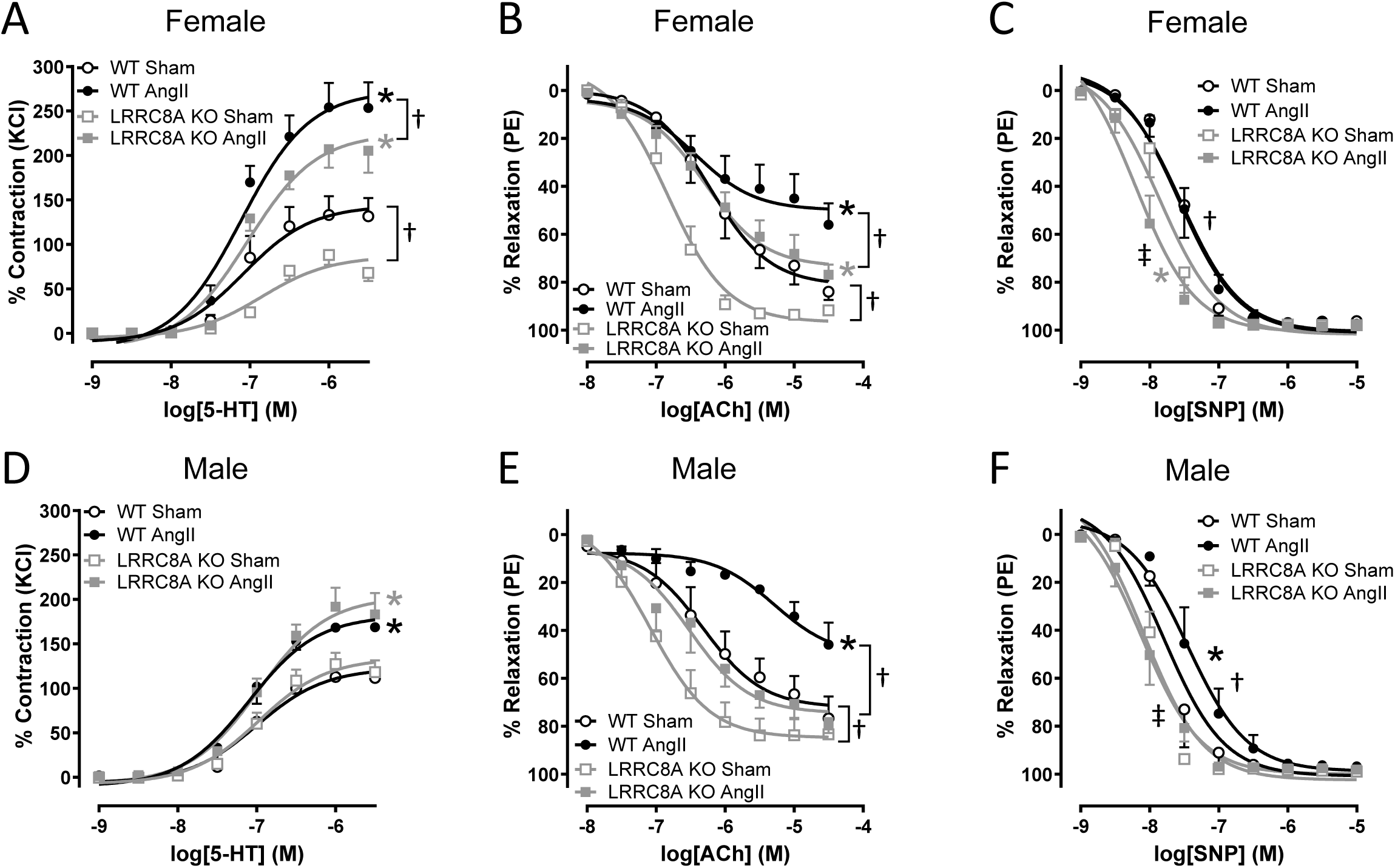
Lack of LRRC8A reduces mesenteric artery dysfunction in female and male AngII hypertensive mice. (**A** and **D**) The maximal contractile response to serotonin (5-HT) was greater in all rings from AngII compared to Sham mice, however only responses of female rings (**A**) were reduced in KO vessels. * Indicates *p* < 0.05 for Sham vs. AngII in WT (black*) or Sham vs. AngII in KO (gray*) for E_max_. † Indicates *p* < 0.05 for E_max_ (n=3 to 5). (**B** and **E**) Relaxation to acetylcholine (ACh) was improved in KO rings in both female and male sham groups. AngII impaired relaxation to ACh in all groups but KO vessels still relaxed much better than WT. The response of KO AngII vessels closely resembled those of WT Sham. Black* indicates *p* < 0.05 compared to WT Sham for E_max_ (**B**) or EC_50_ (**E**). Gray* indicates *p* < 0.05 for KO Sham vs. KO AngII (E_max_ or EC_50_). † Indicates *p* < 0.05 for WT Sham vs. KO Sham (EC_50_) or WT AngII vs. KO AngII for E_max_ (**B**) or EC_50_ (**E**). (**C** and **F**) Relaxation to sodium nitroprusside (SNP) was better in KO compared to WT Sham vessels. AngII impaired relaxation in male but not in female rings, but in both sexes KO relaxed better than WT vessels. * Indicates *p* < 0.05 for WT AngII vs. WT Sham (black*) or KO AngII vs. KO Sham (gray*) for EC_50_. † Indicates *p* < 0.05 for WT AngII vs. KO AngII (EC_50_). ‡ Indicates *p* < 0.05 for WT Sham vs. KO Sham (EC_50_) (n=5 to 7).

Relaxation to SNP was enhanced in KO rings from both male and female mice compared to WT Sham. AngII hypertension impaired relaxation to SNP only in male WT vessels, but KO tissues were protected from this effect. (Fig. 2C and F, LogEC_50_, female WT AngII-7.6 ± 0.07 vs. KO AngII-8.2 ± 0.09; male WT AngII-7.5 ± 0.11 vs. KO AngII-8.1 ± 0.10, \**p*<0.05).

Compared to the responses of mesenteric arteries, aortic rings were less impacted by hypertension and by the loss of LRRC8A. Responsiveness to 5-HT was enhanced by AngII in all groups. In male aortae, contraction of KO rings was reduced in both the Sham and AngII groups compared to WT (Fig. 3A and E, E_max_, male WT AngII 339 ± 12.7 vs. KO AngII 282 ± 6.9%, \**p*<0.05). In contrast, there were no differences between WT and KO responses in females. Contraction to NE was slightly but significantly attenuated in Sham KO aortic rings compared to WT in male mice only (E_max_, male WT Sham 93 ± 5.0 vs. KO Sham 80 ± 3.6%, \**p*<0.05). The response of AngII hypertensive rings to NE was potentiated in all groups but this effect was significantly less in female KO vessels (Fig. 3B and F, E_max_, female WT AngII 159 ± 15.0 vs. KO AngII 111 ± 12.0%, \**p*<0.05). In contrast with mesenteric arteries, aortae showed similar relaxation responses to both ACh and SNP in WT and KO Sham vessels. Relaxation to ACh was attenuated by AngII hypertension in both male and female WT and KO rings, but in females this change was only significant in WT vessels (Fig. 3C and G). The response of WT rings to SNP was mildly but significantly impaired by AngII in both males and females, while KO vessels were not affected (Fig. 3D and H, Log EC_50_, female WT AngII-7.8 ± 0.11 vs. KO AngII,-8.1 ± 0.05, \**p*<0.05; male WT AngII,-8.0 ± 0.07, KO AngII,-8.0 ± 0.07). Collectively these data indicate that vascular dysfunction induced by chronic infusion of AngII and the associated hypertensive milieu was ameliorated in VSMC-specific LRRC8A KO vessels.

**Figure 3.**
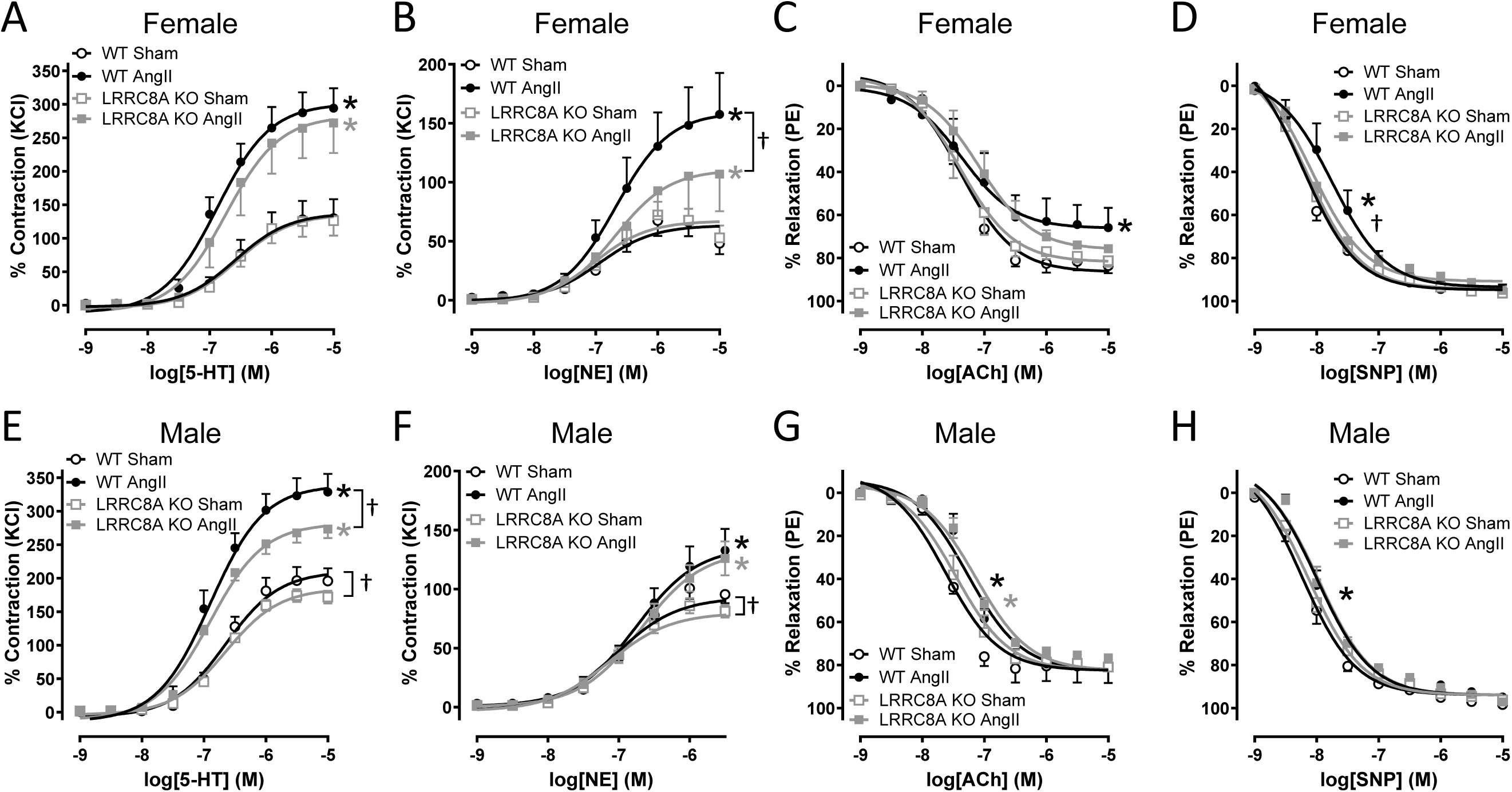
LRRC8A KO reduces dysfunction of aortae from female and male AngII hypertensive mice. AngII-induced vascular dysfunction in aorta was sex-dependent. (**A**, **B**, **E** and **F**) Contractile responses to 5-HT and norepinephrine (NE) were enhanced in all AngII compared to Sham groups. (**A**) In females, contraction to 5-HT was not altered in Sham or AngII KO vessels. (**E**) In males the maximal contraction to 5-HT was reduced in KO compared to WT in both the Sham and AngII groups. (**B**) In females, maximal contraction to NE was unaltered in Sham but lower in KO AngII aortae. (**F**) In males, contraction to NE was significantly reduced in KO compared to WT in Sham but not AngII rings. *Indicates *p* < 0.05 for Sham vs. AngII in WT (black*) or Sham vs. AngII in KO (gray*) for E_max_. † Indicates *p* < 0.05 for E_max_. (**C**, **D**, **G** and **H**) Relaxation to ACh and SNP was slightly more impaired by AngII in WT compared to KO aortae. (**C** and **G**) Maximal relaxation to ACh was reduced in female but not male WT AngII rings and KO responses were unaffected in both sexes. (**G**) In male rings EC_50_ was increased by AngII in vessels from both sexes. (**D** and **H**) In both female and male aortae relaxation to SNP was not altered in Sham tissues but AngII only significantly impaired the response in WT vessels. (C) Black* indicates *p* < 0.05 for WT Sham vs. WT AngII for E_max_. (D, G and H) * Indicates *p* < 0.05 for Sham vs. AngII in WT (black*) or Sham vs. AngII in KO (gray*) for EC_50_. † Indicates *p* < 0.05 for WT AngII vs. KO AngII (EC_50_) (n=3 to 5).

### Phosphorylation of proteins that regulate vascular function

NO^•^ is a critical regulator of vascular reactivity and prime driver of relaxation to both ACh and SNP. It can be inactivated by reacting with O_2_^•-^ to form peroxynitrite. NO^•^ produced by endothelial cells directly turns on soluble guanylate cyclase (sGC) in VSMCs, and cGMP activates protein kinase G (PKG) which phosphorylates and activates the myosin phosphatase target subunit 1 (MYPT1) component of MLCP to promote relaxation. In abdominal aortae the relative abundance of sGCα and β were not altered by LRRC8A KO under Sham conditions.

Both proteins were reduced by AngII and sGCα expression was significantly less reduced in KO compared to WT AngII-infused aortae (Fig. 4). This could contribute to preservation of ACh and SNP relaxation in KO vessels of AngII hypertension.

**Figure 4.**
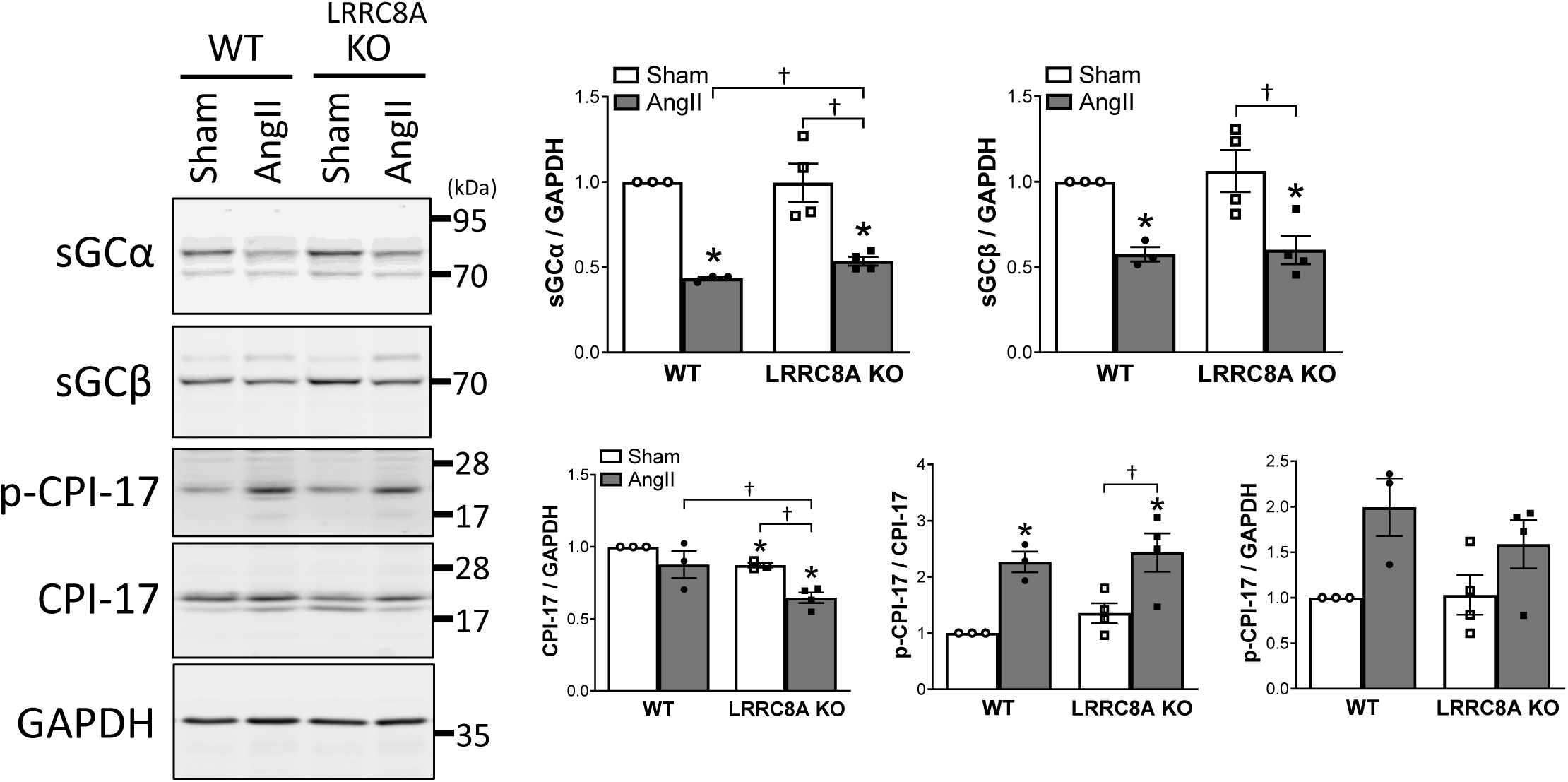
Protein expression related to vascular reactivity is altered in LRRC8A KO aortae. **Top**) Soluble guanylate cyclase subunit (sGCα and sGCβ) abundance was reduced by AngII hypertension in both WT and LRRC8A KO but expression remained significantly higher for sGCα in AngII-treated KO vessels. **Bottom**) CPI-17 protein expression was significantly lower in KO Sham aortae and decreased more in KO AngII treated vessels compared to WT. Activation of CPI-17 (p-CPI-17) was enhanced by AngII in both WT and KO vessels. Bar graphs show the relative abundance of each protein normalized to GAPDH or p-CPI-17 after normalization to total CPI-17. Results are presented as mean ± SEM after normalization to WT Sham. \**p* < 0.05 compared to WT Sham. †*p* < 0.05 (n=3-4).

ROCK and PKC regulate cytoskeletal contractility, motility and stiffness. We previously linked LRRC8A to ROCK-dependent MYPT1 phosphorylation^12^. CPI-17 (protein phosphatase 1 regulatory subunit 14) is a phosphorylation-dependent inhibitor of MLCP and when phosphorylated at Thr38 by either ROCK or PKC it binds to the active site of MLCP, impairs activity, and promotes contraction^28,29^. Total CPI-17 expression was lower in Sham KO compared to WT vessels and this difference was accentuated by AngII. AngII increased CPI-17 phosphorylation (p-CPI-17) to a similar degree in WT and KO vessels (Fig. 4) but decreased CPI-17 abundance may contribute to reduced contractility in LRRC8A KO vessels.

LRRC8A is highly associated with cytoskeletal regulatory proteins and with actin itself^12^. ROCK phosphorylates several of these including ERM and LIMK (Lin-11/Isl-1/Mec-3 Kinase)^30^. ERM proteins link the plasma membrane with actin and phosphorylation increases contractility^31,32^. Total ERM protein in abdominal aortae was not different between WT vs. LRRC8A KO under either Sham or AngII-infused conditions. However, phosphorylation (p-ERM) was reduced in KO compared to WT aortae under both control and AngII-exposed conditions (Fig. 5A). Total ERM was also not altered in cultured mesenteric VSMCs from KO mice, but phosphorylation (p-ERM) was again lower (Fig. 5B). LIMK is activated by ROCK phosphorylation. It subsequently phosphorylates and inactivates cofilin, an actin depolymerizing protein, resulting in actin filament stabilization^30^. LIMK inhibition reduces hypertension-induced arterial stiffening and remodeling^33^. We measured total and p-cofilin to indirectly assess LIMK signaling. Total aortic cofilin was unaltered, but phosphorylation was enhanced by hypertension in WT but not in KO vessels (Fig. 5A). Total cofilin and p-cofilin expression (normalized to tubulin) were both reduced in KO compared to WT in cultured mesenteric VSMCs (Fig. 5B). Finally, phosphorylation of MYPT1 and cofilin was evaluated in cultured VSMCs exposed to

**Figure 5.**
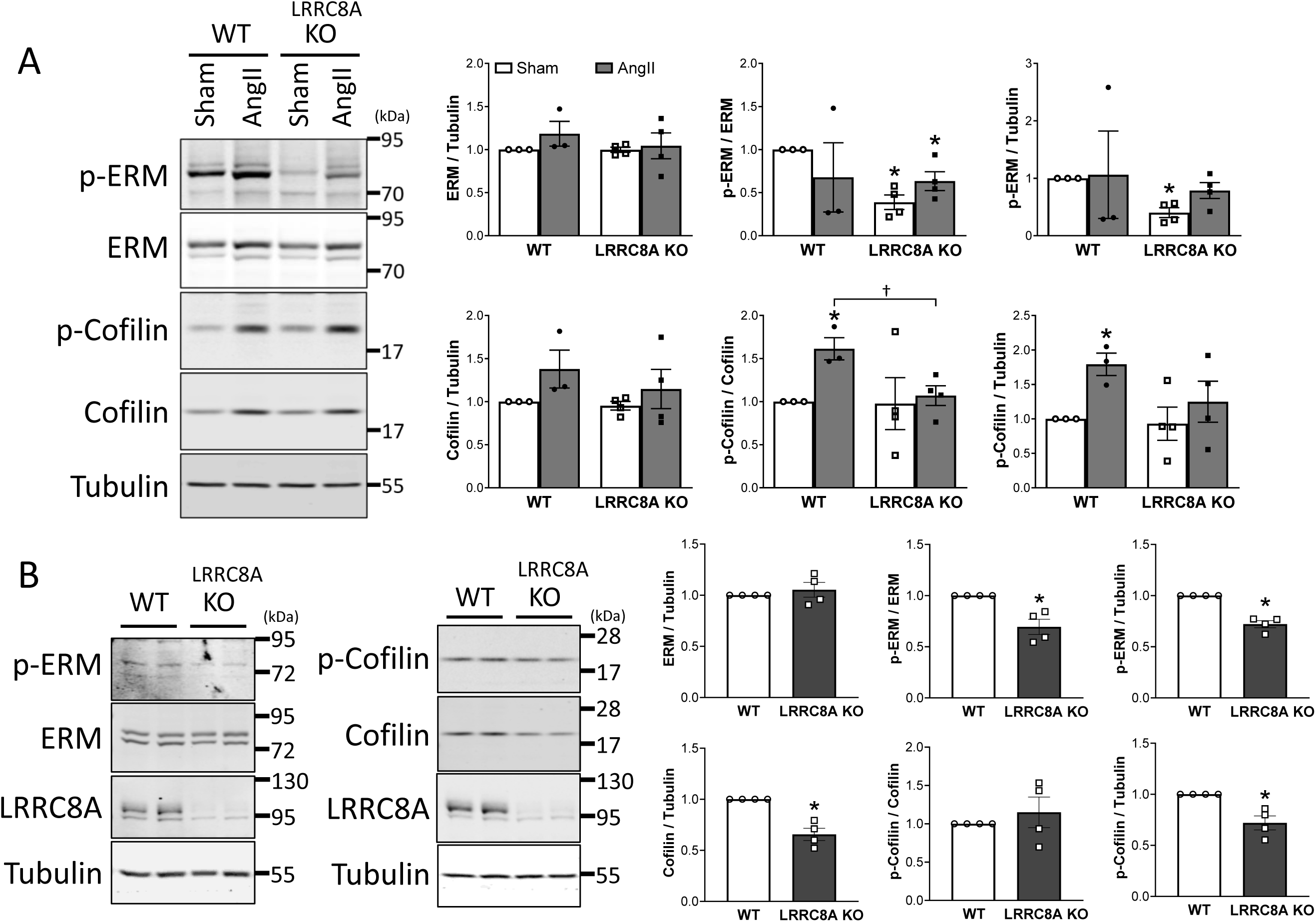
ERM and cofilin expression and phosphorylation in aortae and in cultured mesenteric VSMCs. (**A**) In the aorta total ERM was not altered. Phosphorylation (p-ERM) was not changed by AngII but was reduced in KO compared to WT vessels. Cofilin expression was also not different but phosphorylation (p-cofilin) was increased by AngII in WT but not in KO vessels. \**p* < 0.05 compared to WT Sham. †*p* < 0.05 (n=3 to 4). (**B**) In mesenteric VSMCs, total ERM was not altered but p-ERM was lower in KO cells. Total cofilin was reduced in KO cells and although p-cofilin/cofilin was not different, the total abundance of p-cofilin (p-cofilin/tubulin) was significantly lower in KO cells. Data are mean ± SEM after normalization to WT. \**p* < 0.05 (n=4).

AngII for 3 minutes (Suppl. Fig. S2). AngII significantly increased phosphorylation of both proteins in WT but not in KO cells. These data demonstrate impaired activation of multiple targets of ROCK in LRRC8A KO compared to WT vessels and VSMCs.

### Oxidative stress, inflammation, proliferation and senescence

To assess oxidative stress, we quantified antioxidant enzymes in the aorta (Fig. 6A). As previously observed^34^ cytoplasmic copper-zinc superoxide dismutase (CuZnSOD) expression was not affected by AngII infusion in WT mice and was also not altered by LRRC8A KO. The abundance of extracellular SOD (ecSOD) was similar in WT and KO Sham aortae and was increased in both genotypes by AngII but this change was significant only in KO mice.

**Figure 6.**
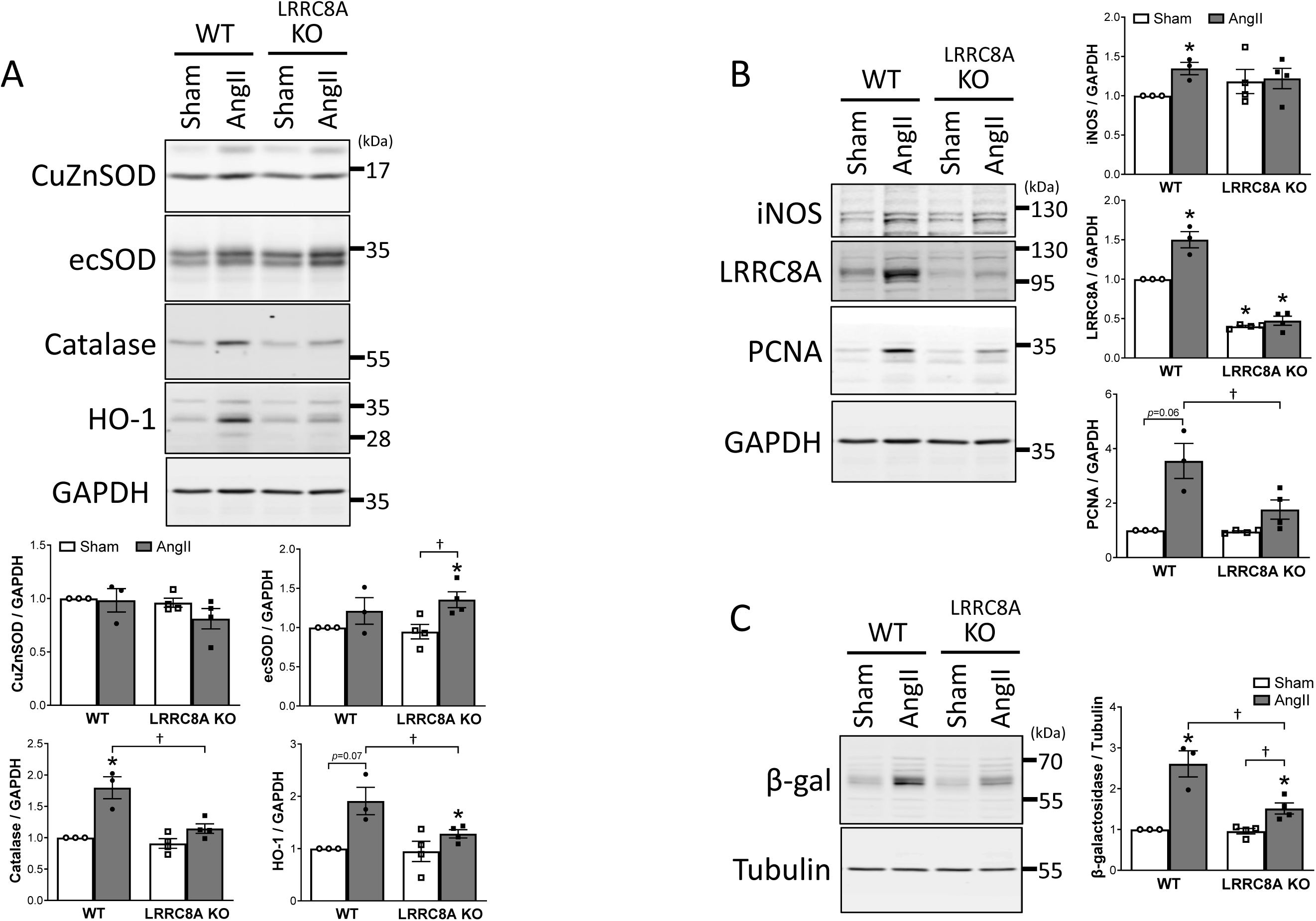
Antioxidant enzyme expression, proliferation and senescence are altered in abdominal aortae from LRRC8A KO mice. (**A**) CuZnSOD expression was not altered by LRRC8A KO or by AngII hypertension. ecSOD was increased by AngII only in KO vessels. Catalase and HO-1 expression were reduced in KO AngII hypertension compared to WT AngII. (**B**) iNOS and LRRC8A expression was enhanced by AngII hypertension in WT aortae. PCNA expression was increased in AngII hypertension (*p* = 0.06), but this effect was reduced in KO vessels. (**C**) The senescence marker β-galactosidase (β-gal) increased by AngII hypertension in both WT and KO, but this effect was reduced in KO vessels. Data are mean ± SEM normalized to WT Sham. \**p* < 0.05 compared to WT Sham. †*p* < 0.05 (n=3 to 4).

Expression of catalase and HO-1 were not altered by LRRC8A KO under control conditions, but both enzymes were induced by AngII hypertension only in WT mice.

Inflammation increases expression of iNOS in VSMCs^35^ and while this effect did not differ in Sham vessels, iNOS was increased significantly by AngII in WT but not KO mice (Fig. 6B). VSMC expression of LRRC8A itself was increased in cerebral vessels from AngII hypertensive mice, and by AngII in cultured VSMCs^17^. In accordance with those prior findings, LRRC8A expression was increased in AngII hypertensive WT mice (Fig. 6B). PCNA expression increases in proliferating cells and while expression was not different in Sham aortae, it increased in Ang-exposed WT vessels (*p* = 0.06) and become significantly higher than in AngII KO mice (Fig. 6B). AngII hypertension is associated with aortic hypertrophy, and in view of the above findings it was surprising that wall/lumen did not differ significantly between aortae from WT and KO AngII hypertensive mice (Suppl. Fig. S3). Beta-galactosidase is a protein marker of cellular senescence^36^. The abundance of β-galactosidase (β-gal) increased more than 2-fold in the aorta of WT mice exposed to AngII but was significantly less induced in aortae from LRRC8A KO mice demonstrating protection from senescence.

## Discussion

VSMC-specific LRRC8A KO augmented BP dipping during inactive periods under control conditions and preserved dipping in the setting of AngII hypertension. Loss of LRRC8A did not alter acute AngII-induced vasoconstriction but vasomotor function was remarkably preserved in hypertensive KO mice. This manifested as reduced contraction and enhanced relaxation and was associated with increased sGCα and reduced CPI-17 protein levels. Our prior finding of reduced RhoA-activity in knockdown VSMCs^12^ was supported by decreased phosphorylation of EMR and cofilin in hypertensive KO vessels. This indicates altered cytoskeletal regulation that may reduce vascular stiffness. Both stiffness and endothelial function are associated with nocturnal BP dipping in humans^37^. Decreased catalase and HO-1 expression in KO AngII aortae suggested reduced oxidative stress, which is associated with reduced senescence^5^ as reflected here by lower expression of iNOS and β-galactosidase.

AngII infusion triggers coordinated cardiovascular, renal, neural and hormonal changes that increase BP and remodel the vasculature. It acts centrally to increase sympathetic drive and vasopressin release, and targeted lesions of the central nervous system dramatically ameliorate AngII hypertension^38^. It also acts on the kidneys to alter afferent and efferent arteriolar function, modifying ultrafiltration and fluid balance, and on the adrenal gland to increase aldosterone secretion which promotes renal sodium retention and vascular inflammation^39^. Here, only VSMC expression of LRRC8A was disrupted thus focusing attention on direct and indirect effects specifically on the vasculature. AngII causes VSMC contraction via influx of extracellular Ca^2+^, mobilization of intracellular Ca^2+^ stores, and increasing the Ca^2+^ sensitivity of the contractile proteins via RhoA/ROCK^40^. AngII also directly activates MAPK and NF-κB signaling to cause endothelial and VSMC inflammation and senescence^41^. Virtually all these effects require production of O_2_^•-^ by Nox1 as a critical element of the signaling cascade^42^. This was demonstrated by the impact of Nox1 knockout on vascular responses to AngII^26^. Loss of regulation of ROS production and oxidative stress are fundamental characteristics of hypertension, but their influence is complex, as demonstrated by the mixed results of clinical trials of antioxidant therapy^10^.

Lower SBP in LRRC8A KO mice was observed only during light (inactive) periods. It is normal for SBP to fall during sleep, a phenomenon that in humans is termed “nocturnal dipping”. Loss of dipping is an independent risk factor for cardiovascular complications of hypertension^43^. It is associated with increased arterial stiffness^44^ and with multiple inflammatory diseases that promote hypertension including diabetes^45^, rheumatoid arthritis^46^, inflammatory bowel disease^47^, lupus nephritis^48^ and chronic kidney disease^49^. Impaired dipping is also correlated with elevation of serum markers of vascular inflammation^45,50^ and endothelial dysfunction^51^. We observed enhanced SBP dipping in LRRC8A KO mice under control conditions, and this difference was amplified by AngII infusion as KO mice retained normal dipping while it was impaired in WT animals. Thus, reduced vascular inflammation in VSMC-specific LRRC8A KO may have contributed to preserved dipping^52^.

In the setting of a relatively modest difference in SBP the preservation of *in vitro* vascular function in LRRC8A KO mice was remarkable, and as observed previously^12^ most marked with respect to vasodilation of mesenteric resistance vessels. TNFα exposure for 48hr in tissue culture increased contraction to NE and impaired relaxation to both ACh and SNP in WT vessels. KO vessels were protected from these effects, completely so with respect to ACh and SNP^12^. We saw similar effects in AngII hypertension, though protection of vasodilation to Ach was less complete than were responses to SNP, perhaps suggesting that 14 days of exposure to AngII caused more significant endothelial dysfunction than did TNFα exposure *in vitro*.

We previously proposed^12^ that the ability of LRRC8A to modulate vasomotor function is related to association of LRRC8A channels with Myosin Phosphatase Rho Interacting Protein (MPRIP), a scaffolding protein that also binds actin, RhoA, and MYPT1. MYPT1 is the regulatory subunit of MLCP and the target of both inhibition by ROCK and activation by PKG. LRRC8A KO reduced ROCK-dependent MYPT1 phosphorylation, and we speculated this was related to loss of RhoA activation by O_2_^•-^ influx through LRRC8A channels^12^. Here we again observed reduced ROCK-dependent MYPT1 phosphorylation in KO VSMCs stimulated with AngII (Suppl Fig. S2A). We also observed changes in the abundance and phosphorylation of another ROCK target that promotes contractility, the MLCP inhibitory protein CPI-17.

Phosphorylated CPI-17 inhibits MLCP to enhance Ca^2+^ sensitivity and smooth muscle-specific CPI-17 overexpression increased vascular contractility and blood pressure^28^. Total abundance of p-CPI-17 was reduced in KO vessels which would be expected to decrease Ca^2+^ sensitivity and reduce contractility. In contrast, sGC is directly activated by NO^•^ and makes cGMP which in turn stimulates PKG to phosphorylate and activate MLCP. The amount of the α or β subunits of sGC in aortae was not altered in KO under normotensive conditions, but expression of sGCα dropped significantly less in KO than in WT arteries in the setting of AngII hypertension. This decrease is known to occur in AngII hypertension^53^ and the reduced drop in KO vessels may contribute to enhanced relaxation responses to the NO^•^ generated by ACh and SNP.

Mass spectrometry analysis of LRRC8A-associated proteins demonstrated interaction with multiple components of the actin cytoskeleton^12^. LRRC8 channel immunoprecipitation pulls down both ERM proteins and cofilin (data not shown). We therefore assessed phosphorylation of these cytoskeletal targets of ROCK. ERM proteins link the plasma membrane with actin filaments and localize to cell surface structures such as ruffled borders and cell adhesion sites along with RhoA^54^. LRRC8A channels, MPRIP and ROS localize to these same structures^3,12^. ERM proteins directly interact with Rho GDI (Rho GDP dissociation inhibitor), activate Rho subfamily proteins, and reorganize actin filaments^55^. ERM phosphorylation was decreased in LRRC8A KO aortae and in cultured KO VSMCs compared to WT, further supporting reduced ROCK activity, and potentially preventing actin filament rearrangements that impair vascular function. ROCK also activates LIMK which phosphorylates and inactivates the actin filament severing protein cofilin, resulting in F-actin stabilization which increases vascular stiffness in AngII hypertension^33^. Cofilin phosphorylation was increased in aortae from AngII-exposed WT but not KO mice, and cultured KO VSMCs had a lower total abundance of p-cofilin.

VSMC AngII signaling has been linked to LRRC8A by protection from cerebrovascular remodeling in AngII infused LRRC8A KO mice^17–19^. Given the established interdependence of LRRC8A and Nox1 in TNFα signaling^16^, unaltered AngII-induced contraction which is Nox1-dependent^24^, needs to be reconciled with reduced AngII-induced inflammation. Despite normal contractile effects of AngII in KO vessels, we did observe reduced acute (3 min) phosphorylation of MYPT1 and cofilin in cultured KO VSMCs (Suppl. Fig. S2), suggesting that at least some fundamental aspects of AngII signaling are altered. It is possible that despite their shared requirement for Nox1, the fundamental relationship between AngII and TNFα receptors and LRRC8A differs. Type 1 TNFα receptors are part of a multi-protein complex that includes Nox1, its p22phox subunit and LRRC8A^12,16,21^. We have speculated that physical proximity of the channels to Nox1 enables channel activation via Nox-induced local depolarization which in turn allows charge compensation of current flow through the oxidase^56^. This physical relationship may not exist for AngII receptors, thus rendering many aspects of AngII signaling LRRC8A-independent.

So why is inflammation reduced, and function preserved in AngII-exposed LRRC8A KO vessels? It is important to consider the contribution of secondary inflammatory pathways that are stimulated by AngII, including hypertensive wall stress itself which activates MAPKs and NF-κB in VSMCs^57^. Aldosterone increases ROS production and NF-κB in VSMCs^58^, and mineralocorticoid antagonism reduced BP in AngII hypertension^59^. The relationship between aldosterone signaling and LRRRC8A has not been explored. TNFα also contributes to AngII-induced inflammation as TNFα KO mice were protected from AngII hypertension, and BP elevation was restored by concurrent TNFα infusion^11^. Impaired TNFα signaling in VSMCs lacking LRRC8A is associated with reduced O_2_^•-^ production, MAPK (JNK, ERK) phosphorylation and NF-κB activation^16,21^. Disrupted TNFα signaling in LRRC8A KO VSMCs may therefore have contributed to protection from AngII. It is important to consider that longer exposure to AngII may have produced more remarkable BP differences and prevented more significant structural phenotypes, as the negative impact of inflammation and senescence clearly worsens phenotypes over time, as is seen in aging^60^. This may explain the protection from structural changes seen in cerebral vessels of VSMC-specific LRRC8A KO mice exposed to AngII infusion for a longer period of 4 weeks^18^.

VSMCs with reduced LRRC8A expression make less extracellular and endosomal O_2_^•-^^16^.

This might be anticipated to reduce oxidative stress which could directly impact vascular inflammation and function. AngII hypertension did not significantly alter expression of either cytoplasmic SOD (CuZnSOD) or extracellular SOD (ecSOD) in WT aortae but ecSOD increased more in AngII exposed KO vessels. ecSOD expression is induced by AngII^34,61^ and mitigates

AngII hypertension^62^. Increased ecSOD in KO vessels may improve NO^•^ bioavailability but may also reflect this fact as expression is activated by NO^•^ in a cGMP and PKG-dependent manner^63^. A report of increased aortic ecSOD expression in AngII hypertension speculated that this occurred as compensation for oxidative stress^34^. However, ecSOD has complex effects on extracellular redox signals, and we showed previously that ecSOD actually supports pro-inflammatory TNFα signaling while tonically inhibiting α5β1 integrin activation^64^. Since loss of LRRC8A reduces extracellular O_2_^•-^, a teleologic case could also be made for cells attempting to correct for impaired O_2_^•-^ bioavailability by decreasing decomposition of extracellular O_2_^•-^.

Catalase expression is increased by aging^65^, heart failure^66^, and is stimulated by lipid peroxides in VSMCs^67^. Transgenic overexpression of catalase can protect against hypertrophy^68^ and abdominal aneurism formation^69^ but not hypertension in AngII-infused mice. Catalase expression increased significantly in AngII-treated WT but not KO vessels. HO-1 is a detoxification enzyme that is responsible for heme catabolism and is vasculoprotective in hypertension^70,71^. Expression is induced by AngII, and by wall^72^ and oxidative stress^71^. However, the direct effect of AngII in cultured VSMCs is to decrease HO-1 expression^73^. The observed decrease in catalase and HO-1 in KO vessels most likely reflects reduced oxidative stress.

AngII causes Nox1-dependent aortic hypertrophy^9^. Here the wall/lumen ratio of AngII hypertensive LRRC8A KO aortae was not significantly different from WT. However, KO vessels exhibited a reduced abundance of PCNA. This protein correlates with proliferation and increases significantly in AngII hypertension^74^. AngII causes redox stress^75^ and induces VSMC senescence^76^ that Nox1-derived oxidants are critical drivers of this process^6^. Evidence for reduced inflammation and lower oxidative stress in LRRC8A KO vessels included reduced expression of iNOS and β-galactosidase. iNOS expression increases with age, is associated with oxidant stress, and is driven by NF-κB^77,78^. β-galactosidase is the primary marker of senescence and expression is closely linked to oxidative stress and associated with enhanced inflammation and accelerated aging^36^. One potential impact of LRC8A on vascular inflammation that we did not assess is the proposed role of these channels in cyclic GMP-AMP (cGAMP) synthase (cGAS)-stimulator of interferon genes (STING) signaling. LRRC8A/E channels can mediate transfer of cGAMP from the cytoplasm to bystander cells to promote STING-mediated, interferon-dependent inflammation^79^ that can drive inflammation and senescence^80^.

VSMC-specific LRRC8A KO did not alter the vasoconstrictor effects of AngII and the BP phenotype of these mice when challenged with an AngII infusion was subtle, consisting only of preserved BP dipping during inactive periods. However, blood vessels from hypertensive KO mice were remarkably protected from inflammation, oxidative stress and vascular dysfunction. These data highlight the vascular impact of the complex inflammatory milieu induced by this form of hypertension and its relationship to BP. They also support our prior observation that loss of LRRC8A reduces RhoA activity, limits inflammation-induced increases in vasoconstriction, and enhances vasodilation under control conditions and preserves it in the setting of an inflammatory challenge. The known association of LRRC8A with cytoskeletal regulatory elements and the reduced phosphorylation of ERM proteins and cofilin observed in LRRC8A KO vessels suggests that this KO may also protect against cytoskeletal disruption and reduce vascular stiffening. This work provides a glimpse of the potential beneficial impact of this anti-inflammatory intervention over a 2-week period. Future work will explore the long-term impact of abrogating VSMC inflammation and senescence on more chronic forms of vascular disease.

## Acknowledgments

We wish to thank Carlo Malabanan and Vanderbilt Mouse Metabolic Phenotyping Center (Vanderbilt University, Nashville, TN) for providing mouse surgery and recording of blood pressure.

